# Model-based dynamic off-resonance correction for improved accelerated fMRI in awake behaving non-human primates

**DOI:** 10.1101/2021.09.23.461491

**Authors:** Mo Shahdloo, Urs Schüffelgen, Daniel Papp, Karla L. Miller, Mark Chiew

## Abstract

**Purpose:** To estimate dynamic off-resonance due to vigorous body motion in accelerated fMRI of awake behaving non-human primates (NHPs) using the standard EPI 3-line navigator, in order to attenuate the effects of time-varying off-resonance on the reconstruction.

**Methods:** In NHP fMRI the animal’s head is usually head-posted, and the dynamic off-resonance is mainly caused by motion in body parts that are distant from the brain and have low spatial frequency. Hence, off-resonance at each frame can be approximated as a spatially linear perturbation of the off-resonance at a reference frame, and is manifested as a relative linear shift in k-space. Using GRAPPA operators, we estimated these shifts by comparing the 3-line navigator at each time frame with that at the reference frame. Estimated shifts were then used to correct the data at each frame. The proposed method was evaluated in phantom scans, simulations, and *in vivo* data.

**Results:** The proposed method is shown to successfully estimate low-spatial order dynamic off-resonance perturbations, including induced linear off-resonance perturbations in phantoms, and is able to correct retrospectively corrupted data in simulations. Finally, it is shown to reduce ghosting artifacts and geometric distortions by up to 20% in simultaneous multi-slice *in vivo* acquisitions in awake-behaving NHPs.

**Conclusion:** A method is proposed that does not need any sequence modification or extra acquisitions and makes accelerated awake behaving NHP imaging more robust and reliable, reducing the gap between what is possible with NHP protocols and state-of-the-art human imaging.

## 1. Introduction

Nonhuman primates (NHPs) serve as useful models for understanding the human brain, due to the many functional and structural parallels between the two species^1^. However, NHP brain imaging involves unique challenges relative to human neuroimaging. For example, the macaque neocortex is 15x smaller in volume, and is 25% thinner compared to the human brain^2^. These considerable scale differences necessitate imaging smaller voxels in order to achieve similar levels of spatial delineation, that inherently reduces the signal to noise ratio (SNR), which is further reduced when undersampling is used to accelerated data acquisition. As a remedy, simultaneous multi-slice (SMS) imaging^3,4^ using bespoke multi-channel receive coils^5^ have been recently employed to enhance statistical power by retaining the temporal degrees of freedom while accelerating the scans in anaesthetised NHPs^6,7^.

Echo-planar imaging (EPI) in awake behaving NHPs is further complicated due to vigorous motion in the behaving animal’s body, hands, jaw, and facial musculature^8^. These movements that inevitably occur, even after mechanical head stabilisation, induce unpredictable *B*_0_ off-resonance changes that are orders of magnitude larger than those induced by respiration or cardiac activity in anaesthetised animals^9,10^. While geometric distortion due to these dynamic effects can be partly addressed in image-based post-processing^11^, off-resonance can render the acquired data inconsistent with the reference calibration data used for unaliasing reconstruction^12^, resulting in additional substantial Nyquist ghosting and residual aliasing artifacts that degrade image quality and potentially obscure the signals of interest. Unlike correcting for distortions caused by static field inhomogeneity, these dynamic reconstruction artifacts are not addressable using image-based post-processing or global ghost correction techniques (Fig. 1), for a number of reasons. Since the significant off-resonant effects result in images with considerable ghosting and aliasing, reconstructed images themselves cannot be used to generate reliable field maps using phase- or magnitude-based methods. Furthermore, methods based on interleaved blip up/down phase-encoding directions cannot be used to estimate dynamic field-maps since it is not guaranteed that the off-resonance is the same between readouts in presence of vigorous motion in NHPs^13^. Hence, accurate estimation of dynamic off-resonance prior to image reconstruction is crucial for high-quality accelerated fMRI imaging of awake behaving NHPs.

**Figure 1:**
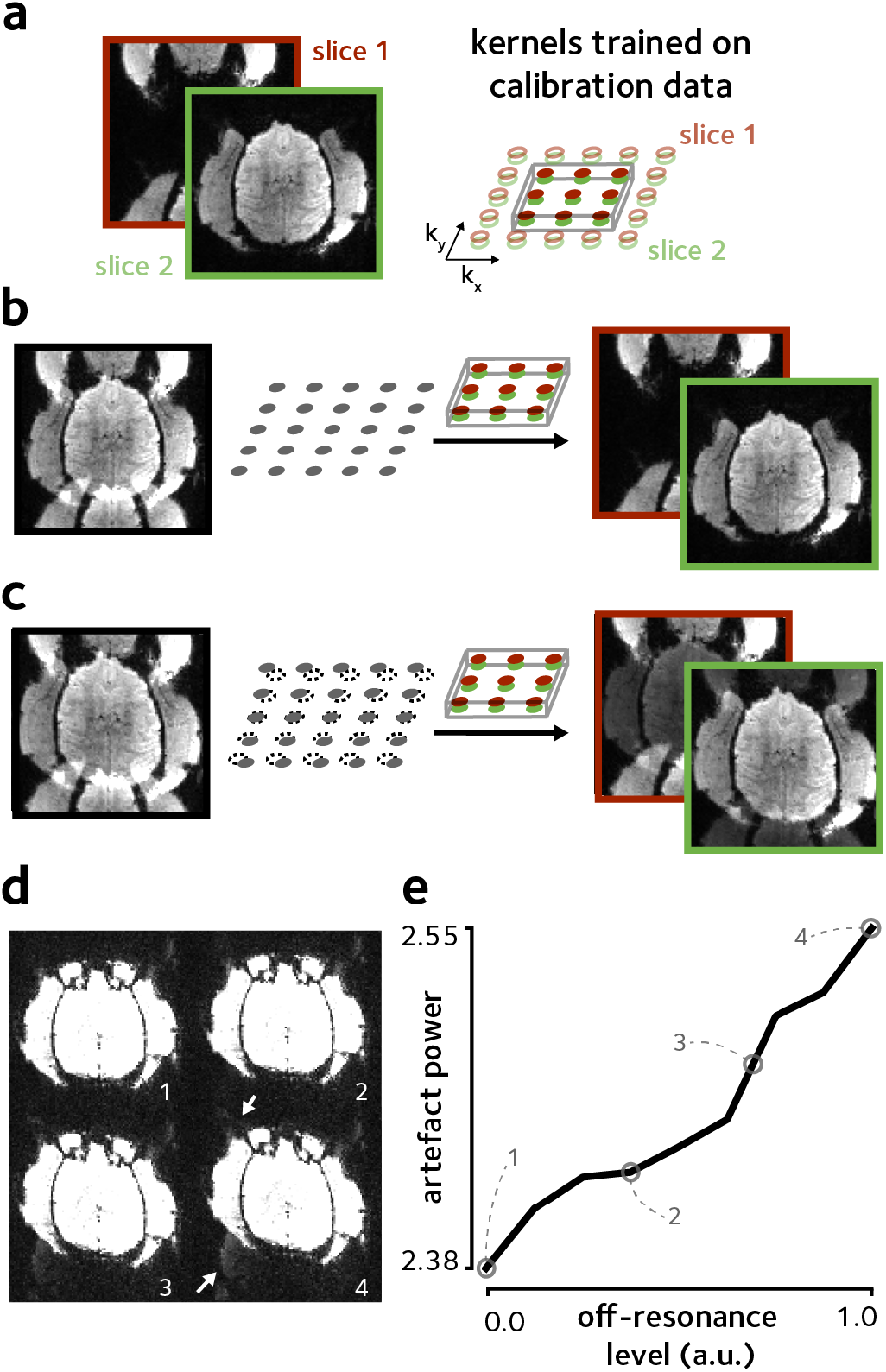
Ghosting artifacts and geometric distortion due to off-resonance. Simultaneous multi-slice accelerated data (MB=2, with CAIPI shifts^30^) were simulated using a single data-frame from fully-sampled *in vivo* macaque data. Simulated spatially linear off-resonance was added. **(a)** The fully-sampled data was used to train split-slice GRAPPA^31^ kernels. **(b)** In the absence of off-resonance perturbations, these trained kernels could be used to completely separate the aliased slices. **(c)** Off-resonance perturbation, however, results in inconsistency of the data to the trained kernels which leads to aliasing artifacts manifested as Nyquist ghosts as well as geometric distortion. **(d)** Increasing the off-resonance level (indicated by numbers) enhances the level of ghosting artifacts and geometric distortion. Here, the display range is saturated to enhance the visibility of artifacts (indicated by white arrows). **(e)** artifact power, taken as the ℓ_2_ norm of the background values, versus off-resonance level. Values corresponding to off-resonance levels in **(d)** are marked by empty circles. artifact power monotonically increases by increasing the levels of off-resonance.

Several approaches have been proposed to estimate dynamic off-resonance in human neuroimaging using field sensors^14–16^, multi-echo sequences^17–20^, or extra navigators^21–24^. In the scope of awake behaving NHP fMRI, these previous works are limited by requiring complicated additional hardware, compromising temporal resolution to accommodate multiple echoes, or requiring customised sequences. Alternatively, a few recent studies have used the spatial encoding provided by multi-channel EPI data, and forward models trained on reference scans or prior information about the tissue structure to estimate dynamic off-resonance in human fMRI^25–27^. Specifically, Wallace et al.^27^ used multi-channel free induction decay navigators followed by calibration of the navigator phase changes using a separate reference scan to estimate and correct the dynamic geometric distortion. Although this study successfully demonstrated geometric distortion correction, it is limited by requiring a separately acquired contrast-matched reference scan and sequence modification.

To mitigate these issues, we propose to use the standard EPI 3-line navigator in order to estimate dynamic *B*_0_ off-resonance changes, without the need for any extra scans, sequence modification, or prior knowledge. This short navigator that samples the central line of k-space three times at each shot is already present in typical EPI sequences and is widely used to correct the Nyquist ghosts caused by discrepancies between odd and even EPI lines. Here we used the generalised autocalibrating partial parallel acquisition (GRAPPA) operator formalism and the spatio-temporal encoding provided by multi-channel 3-line navigator to estimate spatially varying off-resonance at every TR. Dynamic off-resonance estimates were then used to correct the k-space data at each TR in order to make it more consistent with the calibration data, effectively reducing the dynamic ghosting artifacts and also reducing geometric distortions as a byproduct. Performance of the proposed method is demonstrated by successfully estimating the manually introduced off-resonance in phantom experiments, and improving reconstruction quality in simulated and *in vivo* SMS accelerated NHP fMRI.

## 2. Methods

Our proposed method is based on estimating linear k-space shifts in the EPI reference navigator data using GRAPPA operators. In the following sections we first briefly review the concept of GRAPPA operators and their utility for generating arbitrary k-space shifts, and then explain how it can be used in dynamic off-resonance estimation and correction.

### 2.1. GRAPPA operators for k-space shift

In GRAPPA, missing k-space values are synthesised as a linear combination of acquired neighbouring k-space values from all channels, where the neighbourhood used in the linear combination is determined by the geometry of the chosen interpolation kernel^28^. Let 𝒮(**k**) ∈ ℂ^*J*×*N*^ be the acquired data at some k-space location **k** = (*k*_*x*_, *k*_*y*_, *k*_*z*_), *J* be the number of receive channels, and *N* be the number of reconstructed k-space points. In its simplest form, the GRAPPA reconstruction problem synthesizes each missing k-space point using one acquired k-space point from the adjacent phase-encoding line, and can be cast as^29^

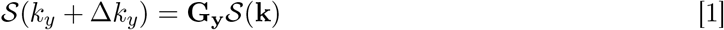

where **G**_**y**_ ∈ ℂ^*J* ×*J*^ is the matrix of GRAPPA weights obtained by calibration on fully sampled data. In this formalism, **G**_**y**_ that maps each k-space point to its adjacent point can be interpreted as an operator that shifts the k-space data one location along the phase-encoding direction. Recursive application of Eq. 1 can be used to produce shifts by multiples of Δ*k*_*y*_

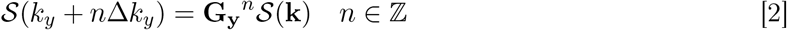

arbitrarily small shifts can be achieved using fractional powers of the operator^32^

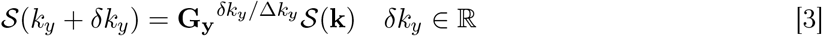

and shifts across multiple directions can be achieved by successive application of operators that were trained on different directions

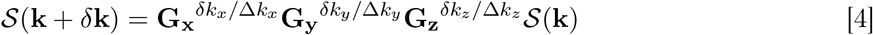

where **G**_**x**_, **G**_**y**_, and **G**_**z**_ are the operators trained to perform shifts of one k-space location along the read-out, phase-encoding, and slice-encoding directions. Here, we trained GRAPPA operators **G**_**x**_, **G**_**y**_, and **G**_**z**_ using the fully-sampled calibration data and used them throughout the reconstruction. Note that the operator in the slice-encoding direction is realisable only with 3D k-space information. To address this issue in SMS EPI, we constructed a proxy 3D calibration dataset that contained information limited to the encoded slices.

### 2.2. Dynamic off-resonance estimation and correction using EPI 3-line navigator

We used the fractional shift property of GRAPPA operators (Eq. 4) and the three EPI navigator lines to estimate dynamic *B*_0_ off-resonance changes at each time-frame of the fMRI time-series. In awake NHP fMRI the head is mechanically fixed but motion in body parts that are distant from the brain causes low spatial frequency off-resonance changes^10^. Therefore, assuming Δ*ω*^*p*^ is the off-resonance frequency during the acquisition of navigator lines at frame *p*, a first order approximation can be used to model that as dynamic perturbations around the off-resonance at the reference frame (*p* = 0)

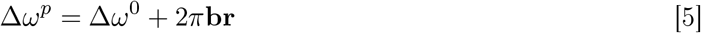

where **b** = (*b*_*x*_, *b*_*y*_, *b*_*z*_) is the vector of coefficients, and **r** = (*x, y, z*)^*T*^ is the spatial location.

Letting 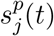 to be the acquired navigator signal from channel *j* at time-frame *p*, the signal in presence of off-resonance can be expressed as

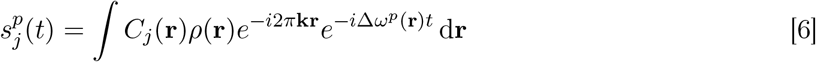

where *C*_*j*_ is the sensitivity profile for channel *j* and *ρ* is the object magnetisation. Using Eq. 5, the navigator signal can be expressed as

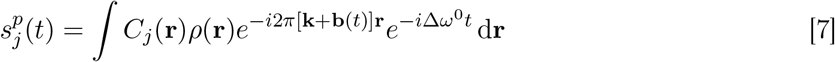

Thus, the navigator data acquired at frame *p* can be modelled as a shifted version of the navigator data at frame 0

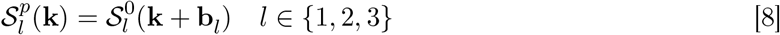

where 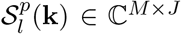 is the *l*th navigator line at frame *p*, and *M* is the total number of k-space columns. Note that because the phase induced by the off-resonance in three consecutive navigator lines varies due to different echo times, the shifts for the three lines are distinct in Eq. 8 (see Fig. 2a). Assuming a constant echo spacing between consecutive EPI line acquisitions the shifts in consecutive navigator lines increase linearly

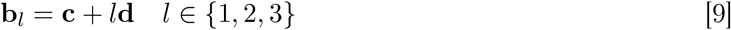

and the first-order off-resonance perturbation at frame *p* can be modelled using the linear shifts in the 3-line navigator data compared to the navigator data at the reference frame

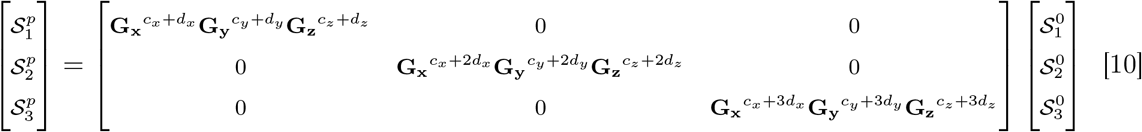

resulting in a model with six unknown off-resonance coefficients and 3*M* × *J* constraints, where **c** = [*c*_*x*_, *c*_*y*_, *c*_*z*_] contribute to the offset off-resonance and **d** = [*d*_*x*_, *d*_*y*_, *d*_*z*_] contribute to the time-varying component of off-resonance.

**Figure 2:**
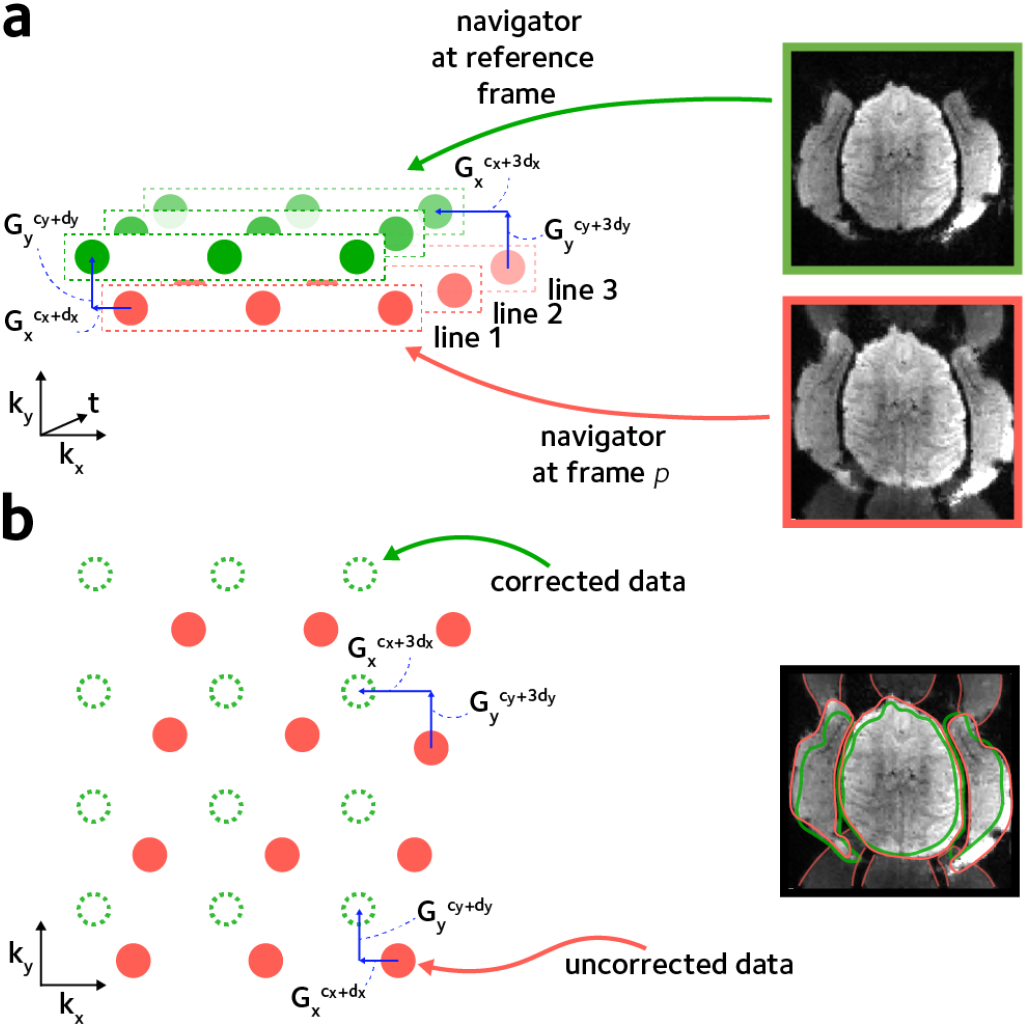
Dynamic off-resonance estimation and correction. Spatially linear off-resonance perturbations can be cast as linear shifts in k-space data. **(a)** To estimate the off-resonance perturbation the 3-line EPI reference navigator data at each TR was compared to the navigator data from a reference frame. Linear shifts between corresponding navigator lines were estimated using GRAPPA operators, accounting for the different echo times of the consecutive navigator lines. **b**. The estimated linear shift coefficients that reflect the off-resonance perturbation were used to correct the EPI data at each TR. This procedure reduces the ghosting artifacts and geometric distortion (red outline), yielding an improved reconstruction (green outline).

The coefficients **c** and **d** at each frame were estimated by solving the problem in Eq. 10 using least squares, and were used to correct the EPI data at each TR. We noted that in frames with unusually high off-resonance the estimated *d*_*y*_ coefficient could be inaccurate. To address such inaccuracies, an additional refinement step was performed. In the refinement step, the estimated coefficients were first used to yield an intermediate image. Then a grid search was performed by perturbing the *d*_*y*_ coefficient around the initial estimate to maximise the product-moment correlation coefficient between the intermediate image and the fully-sampled reference image.

A python implementation of the proposed method is available at https://github.com/shahdloo/nhp_recon.

### 2.3. Experiments

#### 2.3.1. Manual shim manipulation

To validate the performance of the proposed method in estimating dynamic off-resonance, a bottle phantom was scanned on a horizontal 3T scanner using a 15-channel custom NHP receive coil (RAPID Biomedical, Rimpar, Germany) and a gradient-echo (GRE) EPI acquisition with parameters: TE/TR=30/2000ms, FA=90, FOV=192mm, 1.5mm isotropic resolution, 24 slices, multiband acceleration factor MB=2. To mimic dynamic off-resonance changes, separate scans were performed where first-order shim terms were manually adjusted up to ±20*µT/m* in 5*µT/m* increments across acquisitions. The proposed method was then used to estimate the changes in linear shim terms, using the navigator data only.

#### 2.3.2. Dynamic off-resonance correction *in vivo*

To examine the performance of the proposed method in estimating and correcting the dynamic off-resonance effects *in vivo*, 2D fMRI data from an awake behaving Macaque monkey using both in-plane and SMS acceleration were collected. All procedures were conducted under licenses from the UK Home Office in accordance with the UK Animals Act 1986 (Scientific Procedures) and with the European Union guidelines (EU Directive 2010/63/EU). The animal was head-fixed in sphinx position in an MRI-compatible chair (Rogue Research). Data were acquired using a GRE-EPI sequence with parameters: TE/TR=30/2210ms, FA=90, FOV=128mm, 1.25mm isotropic resolution, 40 slices, MB=2, in-plane acceleration factor R=2, 50 frames, using the same 15-channel NHP receive coil. Single-band data, and fully-sampled calibration data were acquired along with the functional data. The single-band data were used to simulate a generic accelerated fMRI data with in-plane and SMS undersampling (MB=2, R=2) corrupted by dynamic off-resonance. Random first-order off-resonance perturbations (0±20Hz, mean±std across the field of view) were assumed at each frame separately for each slice pair, and were used to retrospectively warp the single-band data using multifrequency interpolation^33^. Data were then summed across each slice pair. The proposed method was used to estimate the simulated dynamic off-resonance at each frame, and the estimates were used to correct the data.

Finally, to evaluate the proposed method *in vivo*, dynamic off-resonance was estimated for the whole prospectively undersampled fMRI series, and the data were corrected with the proposed method prior to unaliasing reconstruction using the split-slice GRAPPA algorithm^31^.

### 2.4. Quantitative measures

Shannon’s entropy was used to measure ghosting artifact level in the images, as suggested by previous studies^34,35^. Entropy was calculated as

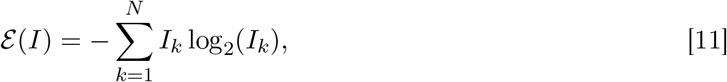

where *I*_*k*_ is pixel magnitude and *N* is the number of image pixels.

To quantify geometric distortion, normalised root mean square error (nRMSE) compared to the reconstructed single-band reference was used. nRMSE was calculated as

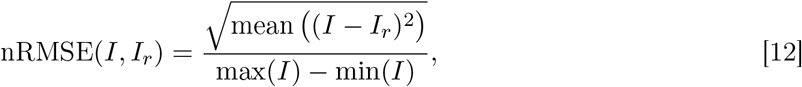

where *I* is the reconstructed image and *I*_*r*_ is the reference image. Note that since the reference image is reconstructed from the undersampled single-band data, the nRMSE measured here would be slightly overestimated relative to an absolute ground truth.

To assess the effect of image quality improvements on the fMRI time series, temporal signal to noise ratio (tSNR) was compared between reconstructions, after masking out the non-brain voxels. While tSNR does not fully characterize the impact of temporally stable ghosting or aliasing artifacts, the metric is useful for capturing dynamic image instabilities that tend to be the main source of ghosting artifacts in awake NHP fMRI. tSNR in pixel *k* was calculated as

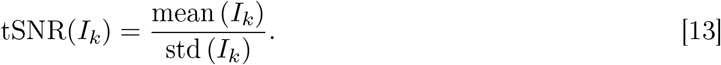

## 3. Results

The performance of the proposed method in estimating manual shim changes in the phantom is shown in Fig. 3. Linear shim term changes in the range of ±20 *µT/m* are accurately estimated with absolute errors of 0.67 ± 0.14 *µT/m* (mean±sem across acquisitions).

**Figure 3:**
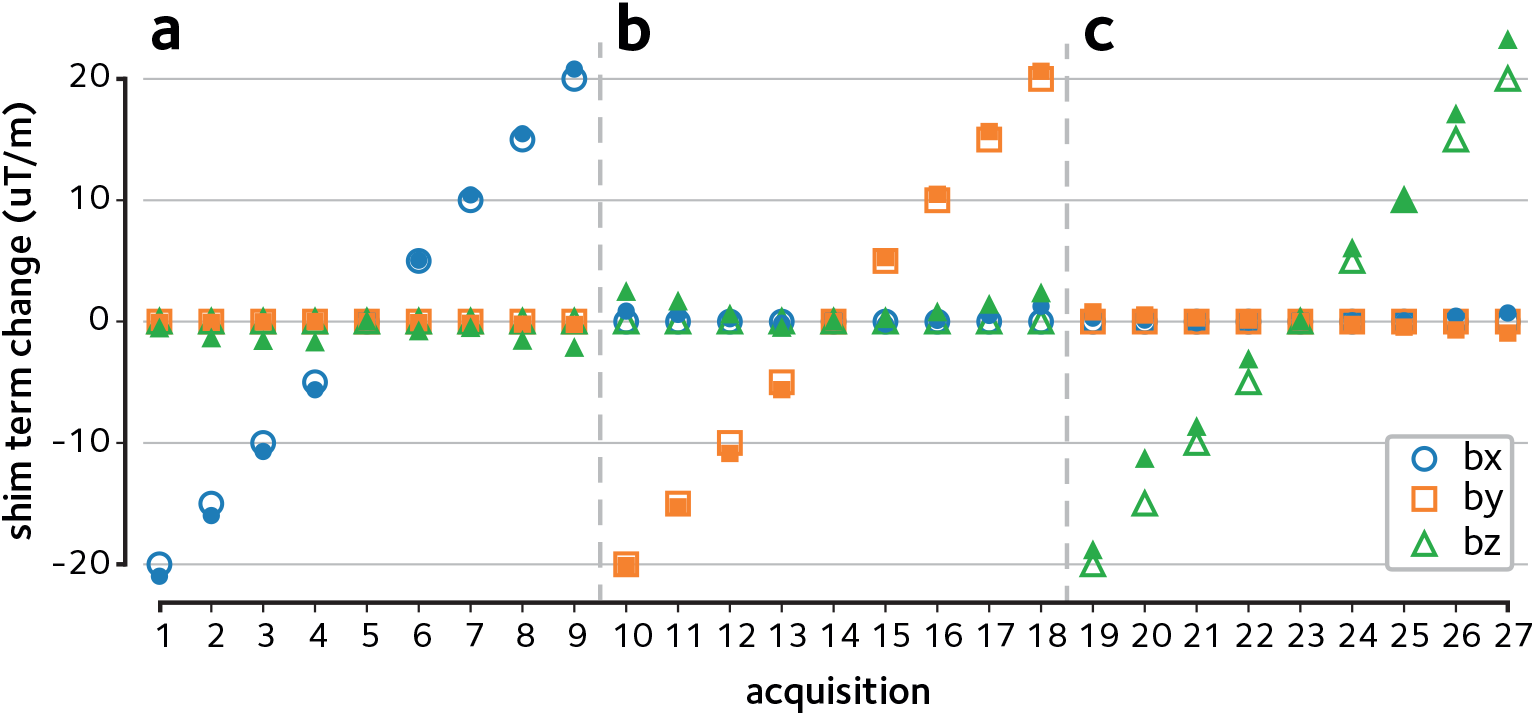
Estimation of linear shim changes in phantom. Dynamic off-resonance changes were induced in a bottle phantom by manually modifying the linear shim terms across acquisitions. The 3-line navigator data were then used to estimate the linear shim term changes across **(a)** x, **(b)** y, and **(c)** z directions in the range ±20*µT/m*. The estimated shim term changes (filled markers) are in good agreement with the ground truth (empty markers).

Next, the proposed method is demonstrated in reconstructions of the *in vivo* data. Figure 4a shows the reconstructions of the retrospectively accelerated data with simulated dynamic off-resonance. Significant ghosting artifacts and geometric distortions that are originally present in the data are successfully attenuated using the proposed approach. Mean image entropy was reduced from 2.19±0.12 *kbits* (mean±sem across time-frames) in the online reconstruction to 1.72±0.10 *kbits* in the reconstruction with the proposed method, a 21% decrease, indicating a significant reduction in ghosting artifacts (bootstrap test, *p* < 10^−4^). Moreover, normalised root mean squared error (nRMSE) compared to the single-band reference image was reduced from 9.43 ± 0.09 % in the online reconstruction to 6.31 ± 0.12 % using the proposed method, indicating a reduction in residual artifacts and geometric distortions.

**Figure 4:**
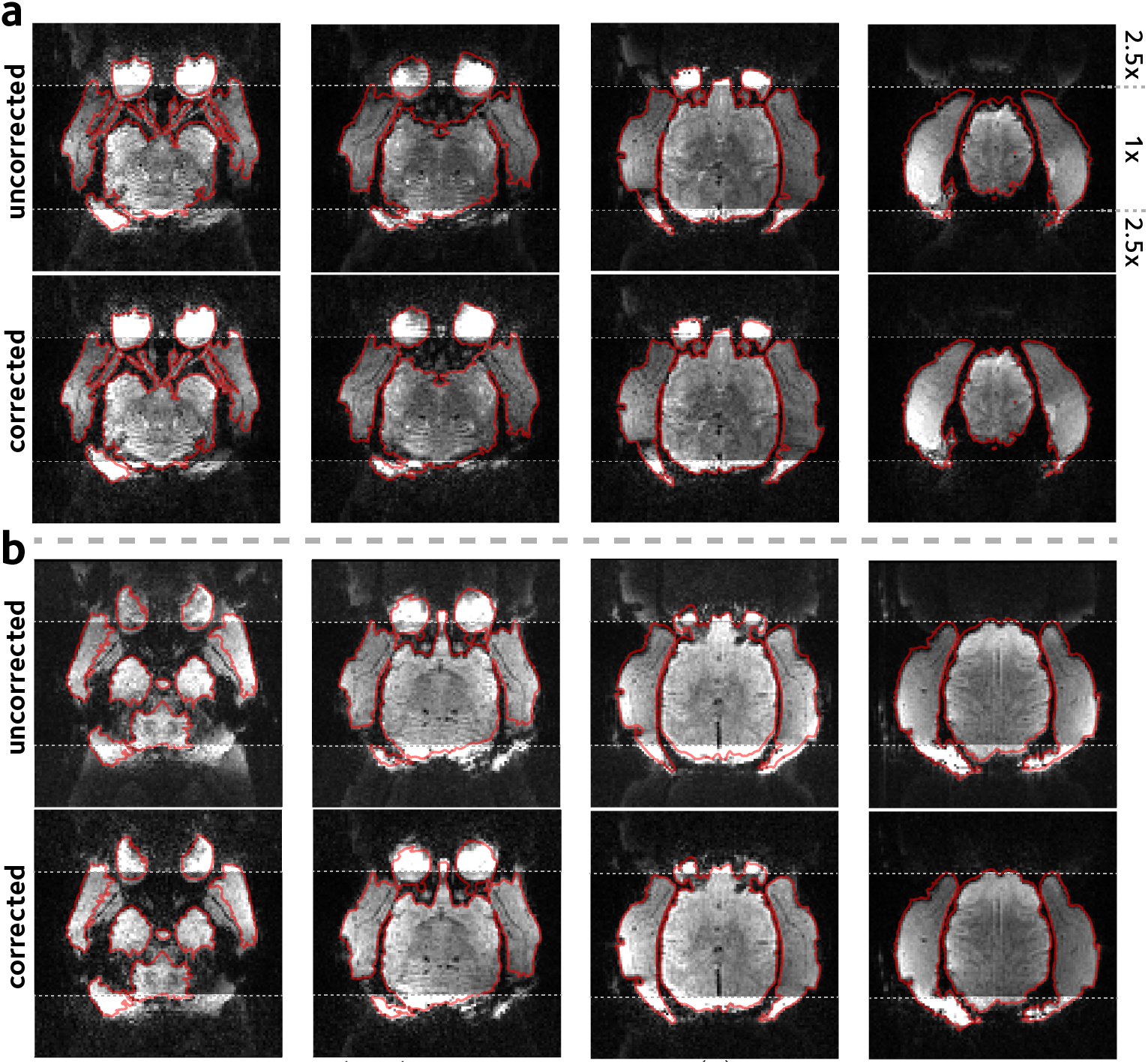
Reconstruction of the *in vivo* EPI acquisition. **(a)** SMS accelerated *in vivo* data were simulated using the single-band k-space data by assuming dynamic off-resonance perturbations. Calibration inconsistency due to off-resonance perturbation causes ghosting artifacts and geometric distortions. 3-line navigator data were used to estimate and correct the simulated dynamic off-resonance. Images from four different slices are shown in columns. The display window in the top and bottom quarter of images are saturated to better show the ghosting artifacts. **(b)** Prospectively SMS accelerated *in vivo* acquisition was corrected and reconstructed using the proposed method. Formatting is identical to panel **(a)**. Estimating and accounting for the dynamic off-resonance yields significantly reduced ghosting artifacts and geometric distortion.

Figure 4b shows a comparison of the proposed reconstruction of the accelerated *in vivo* scan versus the online reconstruction provided as part of the CMRR multiband EPI sequence package^3^, and quantitative comparisons are provided in Fig. 5. Mean entropy was significantly reduced from 2.05 ± 0.01 *kbits* in the online reconstruction to 1.63 ± 0.01 *kbits* in the proposed method (*p* < 10^−4^, Fig. 5a). This 20% decrease is in line with the 21% decrease predicted by the simulations. Normalised RMSE was 8.49 ± 0.04 % in the online reconstruction, and was 6.16 ± 0.02 % in the proposed method (Fig. 5b). These results show that ghosting or residual aliasing artifacts and geometric distortion in the prospectively SMS accelerated *in vivo* scan are considerably attenuated using the proposed method.

**Figure 5:**
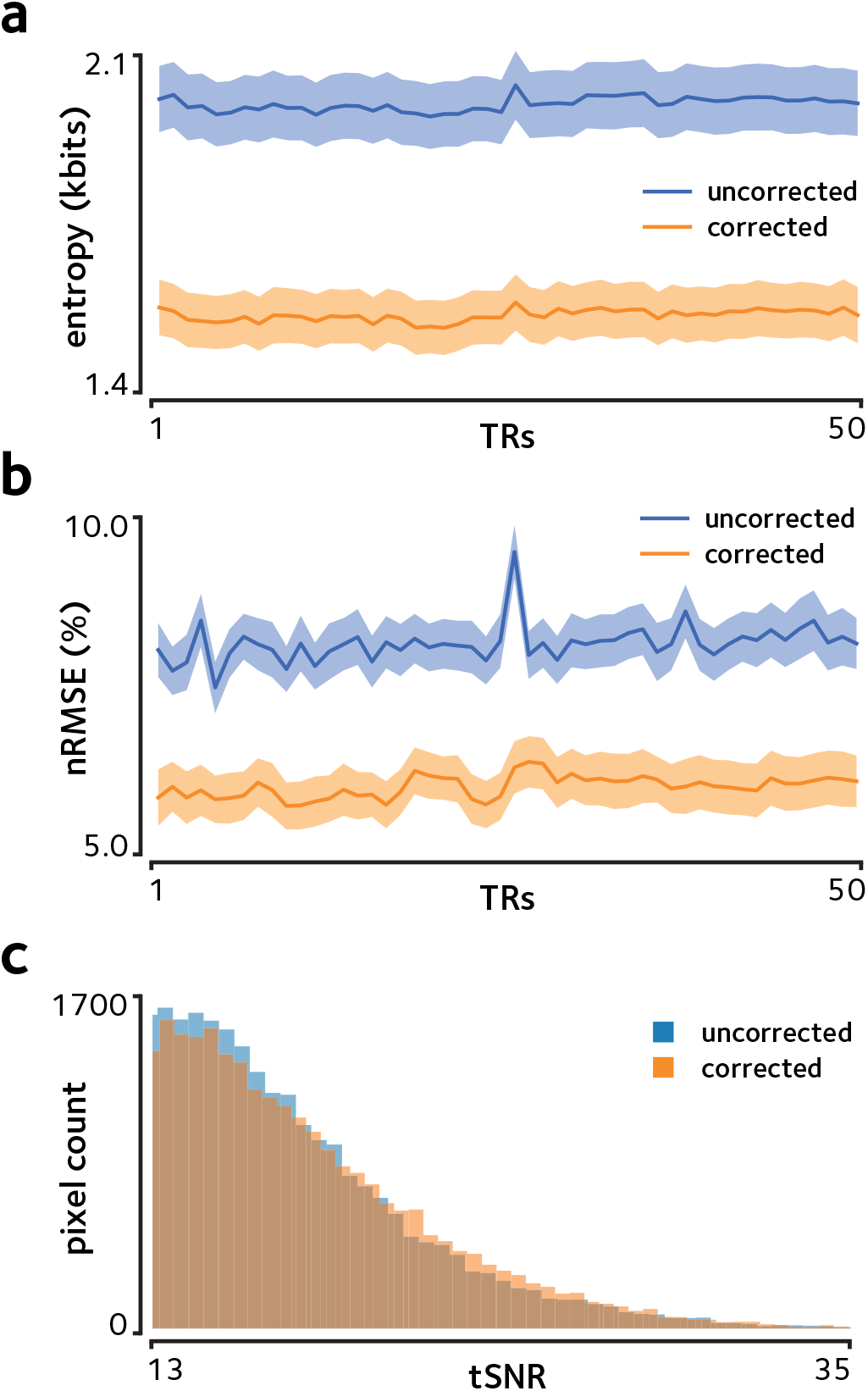
Quantitative measurements for the prospectively SMS accelerated *in vivo* acquisition. **(a)** Image entropy is shown at each TR. Shaded areas show the standard error of the mean across slices. The proposed method decreases the mean entropy, indicating the decrease in ghosting artifacts. **(b)** Normalised root mean squared error compared to the single-band reference image is shown. Shaded areas show the standard error of the mean across slices. Decreased geometric distortion achieved using the proposed method yields reduced nRMSE. **(c)** Histograms of tSNR is compared between the reconstructions. Only the tail of the histogram is shown to compare the distribution in pixels with highest tSNRs. The proposed method yields a histogram that is skewed to the right, indicating higher number of pixels with high tSNR.

Finally, the tSNR was found to be 13.51 ± 0.03 (mean±sem across pixels) in the online reconstruction, and 13.97 ± 0.03 in the proposed method (Fig. 5c), indicating an improvement in functional signal stability.

## 4. Discussion

We have presented a new approach for estimating and correcting the dynamic *B*_0_ off-resonance perturbations in accelerated fMRI of awake behaving nonhuman primates. Acceleration in NHP fMRI has significant impact on data quality and scan efficiency since in-plane acceleration enables achieving higher spatial specificity and SMS acceleration increases statistical significance of the functional time-series. However, vigorous animal motion is unavoidable in imaging of awake and behaving NHPs, even with training. These dynamic effects degrade image quality by making the imaging data inconsistent with the calibration data used during the reconstruction. Our results in phantom, simulation, and *in vivo* experiments show that these effects can be accurately estimated and significantly attenuated using only the standard 3-line EPI reference navigator data. A key property of the proposed method is that it relies only on data acquired using conventional accelerated acquisitions, and does not require sequence modification or lengthened scans to accommodate multiple echoes or more complex navigators. As such, it is also possible to apply the proposed method retrospectively to previously acquired scans, if raw measurement data are available.

In awake NHP imaging, the reduced functional sensitivity is conventionally addressed by using contrast agents^36–39^ or detecting and discarding data frames with excessive motion^40,41^. Using contrast agents biases the functional signal by changing the hemodynamic response functions, hindering direct comparison between findings in NHPs and human studies^42^. Furthermore, detecting and discarding data frames with excessive motion requires extra monitoring hardware and leads to a reduction in the available temporal degrees of freedom. In contrast, the approach presented here can be used to enhance functional data quality in existing NHP acquisition protocols without sacrificing time-points or requiring extra hardware.

While we have focused on demonstrating the proposed method in the context of NHP fMRI, this approach can be applied to other preclinical imaging applications that employ an accelerated EPI acquisition containing the standard 3-line reference navigator, such as accelerated rodent fMRI. Moreover, this approach could have the potential to improve functional sensitivity in human neuroimaging applications in patients with uncontrolled movements such as in Parkinson’s disease, or in neonatal children.

Note that we have have developed the problem with the assumption of having three navigator lines. This results in a model that can be cast as an overdetermined system of equations. However, the model is trivially generalisable to arbitrary number of navigator lines, as long as they are acquired at each TR.

Here, we have assumed first order spatial off-resonance perturbations that have been shown to be a good approximation in imaging headposted NHPs^10^. The theoretical k-space navigator shifting framework in which we cast the problem relies on the first order perturbation assumption to treat the perturbations as additional linear encoding fields. The benefit of this is that we can use existing GRAPPA operator tools to facilitate off-resonance estimation. One limitation of this work, however, is that the accuracy of the first order approximation can suffer with extreme body motion. A potential solution for such extreme cases is to use the proposed approach to provide a near optimal initial guess to be used in a nonlinear image-based dynamic off-resonance correction method, which would be an interesting direction for future research.

In conclusion, the method presented here enables more robust and reliable NHP imaging in accelerated acquisitions, with no sequence or protocol changes, reducing the gap between what is possible with NHP protocols and state-of-the-art human imaging.

## 5. Acknowledgments

The authors would like to thank Steen Moeller for providing the software code used to parse the CMRR MB EPI sequence raw data. M.C. is supported by the Royal Academy of Engineering (RF201617/16/23). The Wellcome Centre for Integrative Neuroimaging is supported by core funding from the Wellcome Trust (203139/Z/16/Z).

